# Chimeric Antigen Receptors Transmit Piconewton Forces that are Coupled with T Cell Function

**DOI:** 10.1101/2025.10.23.684052

**Authors:** Rachel M. Fitzgerald, Areeba A. Hashmi, Anna M. Davis, Alexander K. Foote, Gianna M. Branella, Jhordan Rogers, Emily C. Sullivan, Mohamed Husaini Bin Abdul Rahman, Sunil S. Raikar, Khalid Salaita

**Author notes:** Correspondence should be addressed to Khalid Salaita and Sunil Raikar. **Email:** and.

## Abstract

Chimeric antigen receptor (CAR) T cells are engineered to display a receptor that binds antigens expressed on the surface of cancerous cells, which leads to cancer cytotoxicity. Recently, the T cell field has come to recognize the role of small, piconewton level forces in establishing specificity and cytotoxicity in naïve T cells with the αβ TCR, raising the possibility that these forces could be present in CAR T cells. Using DNA-based tension probes, we reveal 8-19 pN mechanical forces with ∼1 sec timescales transmitted by CAR T cells to their target antigens. CAR-antigen force magnitude is independent of CAR expression level and shows heterogeneity across different T cell donors, suggesting utility as a biomarker of T cell fitness. Using an established exhaustion model, we show strong correlation between CAR exhaustion, cytotoxic capacity, and CAR-antigen force, suggesting that CAR mechanics provide a biomarker of CAR potency complementary to functional assays. Pharmacological inhibition studies demonstrate that CAR forces are driven by actin, Zap70 and Src family proximal kinases. Titration of dasatinib, a clinically used tyrosine kinase inhibitor also dampens both CAR-antigen tension and CAR function in a dose-dependent manner. Structural engineering of the CAR confirms that force levels are modulated by the scFv receptor and co-stimulatory domains, but force transmission requires CD3ζ ITAMs. Taken together, this work shows that CAR T cells transmit pN force to their cognate antigens which holds potential significance in the design and screening of CAR therapeutic candidates and for predicting treatment outcomes in a personalized fashion.

## Introduction

Chimeric antigen receptor (CAR) T-cell therapy, a cancer immunotherapy in which T cells are genetically engineered to target specific cancer surface antigens, has shown great promise in improving outcomes in patients with aggressive cancers.^1, 2^ These successes have primarily occurred in patients with relapsed and/or refractory hematologic malignancies, with FDA approval now for CAR T-cell products targeting B-cell acute lymphoblastic leukemia (B-ALL), large B cell lymphoma, follicular lymphoma, mantle cell lymphoma as well as multiple myeloma.^3^ Despite clinical successes, patient response is variable and a variety of challenges persist. For example, CAR T cell exhaustion, antigen escape and on-target off-tumor effects are all associated with poor clinical outcomes.^4–7^ While there has been tremendous progress, there is still an incomplete understanding of CAR T cell activation and signaling mechanisms. Specifically, the biophysical and mechanical signaling remains understudied, despite being significant for αβ T cell signaling and cytotoxicity.^8, 9^

CARs are often engineered to be expressed on T cells, which naturally express T cell receptors (TCRs). Both receptors mediate activation and cytotoxicity of the T cell but have significant structural differences. Specifically, CARs are single pass transmembrane proteins with an ectodomain presenting a targeting domains such as an single chain variable fragment (scFv) derived from a monoclonal antibody targeting a surface antigen, while the intracellular domain typically includes the cytoplasmic domain of costimulatory CD28 or 4-1BB, and a CD3ζ signaling domain containing several immunoreceptor tyrosine-based activation motifs (ITAMs) (Scheme 1, right). In contrast, the TCR is a complex of multiple transmembrane proteins with an antigen recognition domain composed of an α and β chain heterodimer that associates with intracellular signaling domains, which interact with proximal kinases to drive phosphorylation of three ITAMs on each CD3ζ (Scheme 1, left). The affinity for antigens is also vastly different, with CARs displaying Kd values in the nM range and TCRs showing affinities three orders of magnitudes weaker at μM.

Recent work on the TCR-pMHC complex has shown that the TCR experiences mechanical forces that mediate triggering and tune responses.^8, 9^ Single molecule force spectroscopy and molecular force probe measurements support this model and show that the αβ TCR responds to and transmits 10-20 piconewton (pN) forces to cognate antigens.^10, 11^ A major question in this field relates to whether CARs also transmit forces to their antigens and thus function as mechanotransducers. On one hand, this seems unlikely due to structural differences in the cytoplasmic domains that would hinder cytoskeletal engagement, and lack of a force sensitive FG loop in the extracellular domain that Lang and Reinherz have shown contributes to TCR mechanotransduction.^12, 13^ Moreover, force spectroscopy measurements show that CAR-antigen bonds decrease in lifetime in response to pulling forces (slip bonds), unlike TCR-pMHC bonds, which increase in lifetime under pN level forces (catch bonds).^10, 14^ In native T cells, TCRs cluster to exclude bulky phosphatases, but this clustering and organization is not observed in CARs, potentially limiting this mechanism.^15^

Conversely, CAR-CD19 and TCR-pMHC have similarly sized ectodomains, creating a localized pinch point for membranes. Prior work on T cells and CAR T cells has shown that both are sensitive to changes in the height of this interaction pair.^16, 17^ Interestingly, CD28 stimulation has been suggested to transduce force by increasing TCR traction forces through phosphoinositide 3-kinase (PI3-K) recruitment, but it is unknown if the intracellular domains alone, as displayed in CARs, are sufficient to show similar effects.^18^ The CD3ζ internal domains of the CAR are phosphorylated during activation similarly to an αβ T cell’s endogenous CD3ζ, but the actin and force-bearing proteins are not thought to physically link to the CD3ζ domain.

Due to the structural similarities and clear differences in CARs and TCRs, we became interested in whether a CAR would transmit pN level forces to its cognate antigen. To address this question, we use DNA force probes to measure CAR-antigen forces. Briefly, these probes are comprised of a DNA stem-loop hairpin that presents cell-specific ligands at one terminus and is anchored to a surface through the second terminus (Figure 1A).^19^ When a force greater than the F_1/2_ of the hairpin is exerted on the ligand, the stem-loop is opened, causing a fluorophore-quencher pair to separate and creating a real-time turn-on fluorescence response.^20^ To accumulate transient force events, we utilize a DNA locking strand that selectively binds to the unfolded stem-loop region of the hairpin, preventing the hairpin from refolding (Figure 1F).^21^

**Figure 1.**
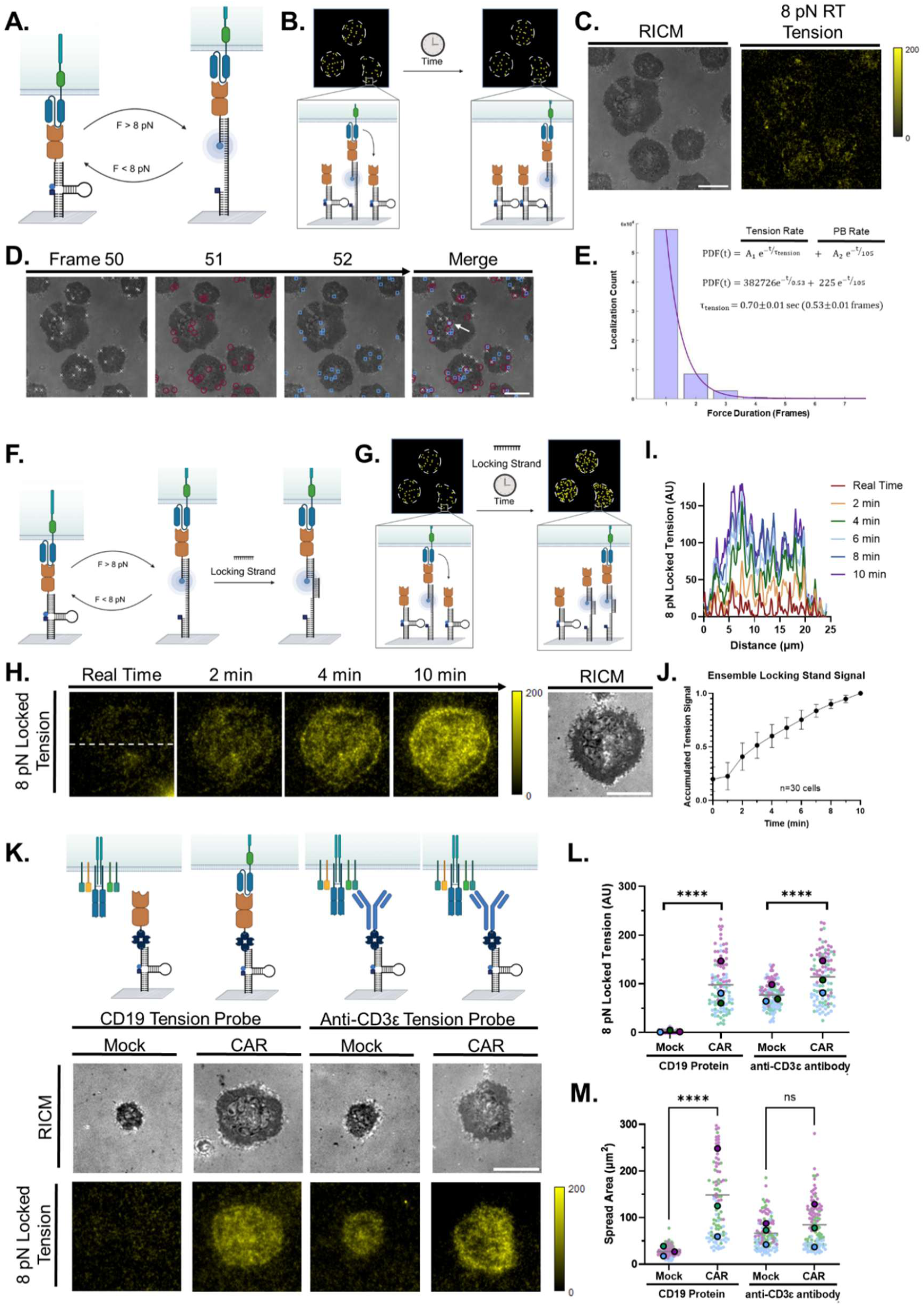
CD19-Directed CAR T Cells exert pN level forces on CD19 antigens. **A** and **B** show schematics of DNA hairpin tension probes reversibly opening and closing in response to dynamic 8 pN forces transmitted through the CAR-antigen complex. **C.** Reflection interference contrast microscopy (RICM) image showing cell-surface contact area, and CD19 CAR 8 pN realtime tension (Cy3B fluorescence) signal. **D**. Overlay of RICM and single molecule localizations of CAR time tension events at *t*= 50 (white), 51 (red) and 52 frames (blue), as well as a merge of all frames showing sustained mechanical events across 3 frames (white arrow). **E**. Histogrammed tension lifetimes (light purple) fit to exponential decay function (dark purple). CD19 CAR mechanical events display τF =0.70±0.01 s. n=87,333 events from 71 unique cells. **F** and **G**. Schematics showing the locking strategy to accumulate mechanical events across time. **H**. Representative Cy3B images showing increasing accumulated CD19 tension signal underneath a single CAR T cell over a 10 min duration after addition of 200 nM locking strand. RICM image was captured at the initial time point and displays the cell contact area. **I.** Line scan intensity analysis of region highlighted by dashed line in H. Colors indicate the time after addition of lock strand. **J.** Ensemble CD19-CAR 8 pN tension signal measured 0-10 minutes after adding locking strand, n=30 cells. The signal was normalized to the maximum integrated intensity at t=10 min. **K**. Representative RICM and 8 pN tension images showing the tension and spread area of cells on CD19 and CD3ε surfaces. Mock cells were not transduced with CAR lentivirus. **L.** Quantification of mean tension per cell, n=90 cells/condition from 3 transductions. Each color indicates a biological replicate from a unique transduction using the same human donor. The larger data points indicate the mean of each replicate while the small data points represent the single cell measurements. **** <0.0001. M. Quantification of cellular spread area across n=90 cells from 3 transductions using the same human donor. **** < 0.0001, ns = 0.3652. All scale bars = 10 microns.

We demonstrate that CD19-directed CAR T cells derived from the FMC63 antibody specifically exert 8-19 pN forces to their antigens with lifetimes at the second time scale. We further show evidence that CAR T cell forces are donor-dependent and are faithful predictors of cytotoxicity *in vitro*. We show that the interactions of internal domains on CAR T cells, including ITAMs and costimulatory domains, have crucial force-transducing roles. Forces are dependent on both cytoskeletal proteins and receptor tyrosine kinases, and inhibition of either of these pathways decreases tension. We finally show that increasing doses of dasatinib, a tyrosine kinase inhibitor, downregulates both CAR tension and cytotoxicity.

## Results

### CAR T Cells Specifically and Rapidly Generate Short-Lived pN Level Forces

As a clinically relevant CAR T cell model, we transduced T cells from healthy human donors with lentiviral vector encoding a second-generation CD19-directed CAR containing a CD28 costimulatory domain and a CD3ζ signaling domain (SI Figure 1). The CD19 binding domain was the scFv derived from the murine monoclonal antibody FMC63, which is currently used in four FDA approved CD19-CAR T-cell products.^22, 23^ To test whether these cells generate forces, we used 8 pN probes displaying CD19 antigen (Figure 1A, 1B). These hairpin probes reversibly open and close in response to force. As soon as cells engaged the surface, we observed punctate, weak and transient fluorescence signal underneath CAR T cells (Figure 1C, SI Video 1, left). This observation confirmed the first evidence that CARs transmit F>8 pN forces to their antigens. Interestingly, the weak real time signal was reminiscent of single molecule events, which was subsequently confirmed based on underlying single-step intensity changes (SI Figure 2E, 2F) and Gaussian shapes (SI Video 2).

To analyze the lifetime of CAR tension events, we imaged at a frequency of 0.75 Hz, background subtracted the resulting signal and identified fluorescent puncta using ThunderSTORM (SI Figure 1, SI Video 1, right). The vast majority of events (>99%) were under the cells and persisted for 1-4 frames (Figure 1D). Force lifetimes were inferred by identifying the persistent puncta that survived across multiple frames (Figure 1D, white arrow shows an example). The distribution of tension-lifetimes was histogrammed and fit to an exponential decay curve, revealing a lower bound of tension lifetime of 0.70±0.01 s (n = 87,333 events) (Figure 1E), much faster than the rate of photobleaching (SI Figure 1). CD19-CAR force lifetimes were consistent with force spectroscopy measurements indicating mean bond lifetimes of 0.4 and 2 sec, for applied forces of 25 pN and 15 pN, respectively. ^14^

Due to the low frequency and rapid interaction of CAR and CD19 in real time, we employed a locking strand which selectively hybridizes to opened force probes, preventing refolding and signal loss once the interaction ends (Figure 1F).^21^ The use of the lock allows for an accumulation of signal over time likely due to multiple CAR engagements with proximal CD19 antigen, resulting in a time-dependent increase in signal (Figure 1G).

After allowing cells to spread on a bed of hairpins displaying CD19 antigen for 10 minutes, 200 nM locking strand was added to each well and imaged over time. Over a 10 min timelapse, the signal intensity increased relative to real-time measurements, confirming that the CAR mediates a large number of short-lived mechanical events (Figure 1I). Representative line scan analysis showed that force events rapidly accumulated at both the periphery of the cell and throughout the cell area (Figure 1H). When we average 8 pN tension signal from n = 30 cells, we found that the signal increased by three-fold and approached steady state at *t*=10 minutes after adding lock (Figure 1J). Importantly, this rate of lock signal accumulation was consistent with the single molecule force lifetime analysis, as low rates of real-time signal and fast accumulation with locking strands implies a short force lifetime. When tracking the history of real-time tension events (SI Video 1, right), the localization of signal mirrors that of cells with locking strand added.

We next aimed to assess the specificity of these events with the use of a mock transduced T cell (mock), which underwent identical processing to the CAR but did not receive a CAR lentiviral construct and cannot recognize the CD19 antigen. When testing both the mock and CAR T cells, both accumulated force events for the anti-CD3ε surfaces, showing mock cells could generate a typical tension response with the TCR complex through CD3ε, as described previously (SI Figure 3, Figure 1K, 1L, 1M).^11^ The tension response to CD19 antigen was markedly different, with CAR cells having 5.22 fold higher spread area and 33.8 fold greater fluorescence signal than mock (****<0.0001, n≥90 cells per condition from 3 independent transductions, Figure 1K, 1L, 1M), showing that the generation of CD19 forces is receptor-dependent. We observed a similar effect using an anti-CD19 CAR antibody, with mock T cells lacking a CD19-directed CAR not spreading or interacting with a CAR-specific antibody (SI Figure 4). These results indicate that CAR T cells can rapidly generate biomechanical forces and that these forces are CAR-specific.

### CARs Transmit F>19 pN at Periphery of Cell-Substrate Junction

Past work from our lab has shown that the OT-1 TCR can transmit *F*>19 pN to its cognate ligand.^11, 24, 25^ Those studies showed rapid accumulation of force events using low threshold probes due to fast kinetics of binding. However, those interactions are catch bonds and the CAR-antigen interaction is a slip bond, raising doubts over this interaction’s capacity to exert greater magnitude of forces. However, single molecule force spectroscopy showed the CD19-CAR complex displayed 12 second and 2 second bond lifetimes with *F* = 10 pN and 15 pN, respectively, and thus we speculated that CAR T cells may transmit greater magnitude forces to the CD19 antigen.^14^

We created greater F_1/2_ CD19 force probes to study this hypothesis. By altering the length and GC content of the stem and loop of the DNA hairpin, we tuned the F_1/2_ of the hairpin from 8 to 19 pN of force (Figure 2A, SI Figure 5). Compared to the more even tension footprint on the 8 pN probes, cells seeded on the 19 pN force probe substrate had a spatial distribution of mechanical events more confined to the cellular periphery, where past work showed enhanced dynamics of actin (Figure 2B).^26–28^ We also observed changes in the number of force events that exceeded the F_1/2_ of the hairpin at the 12 and 19 pN force thresholds. Compared to the 8 pN CD19 probe, this led to a 44.6% decrease in signal with the 12 pN probe and a 69.0% decrease with the 19 pN force probe (Figure 2C). Similar trends were observed for the CD3ε antibody (aCD3ε) probe, with a 39.8% decrease using the 12 pN probe and 88.3% decrease in signal using the 19 pN probe (Figure 2C). Notably, the CD19 19 pN condition generated some level of signal greater than the mock 8 pN tension measurement (P<0.0001), which had no appreciable interaction with a CD19 surface and thus generated no signal, showing a subset of CD19 force events have *F*>19 pN.

**Figure 2.**
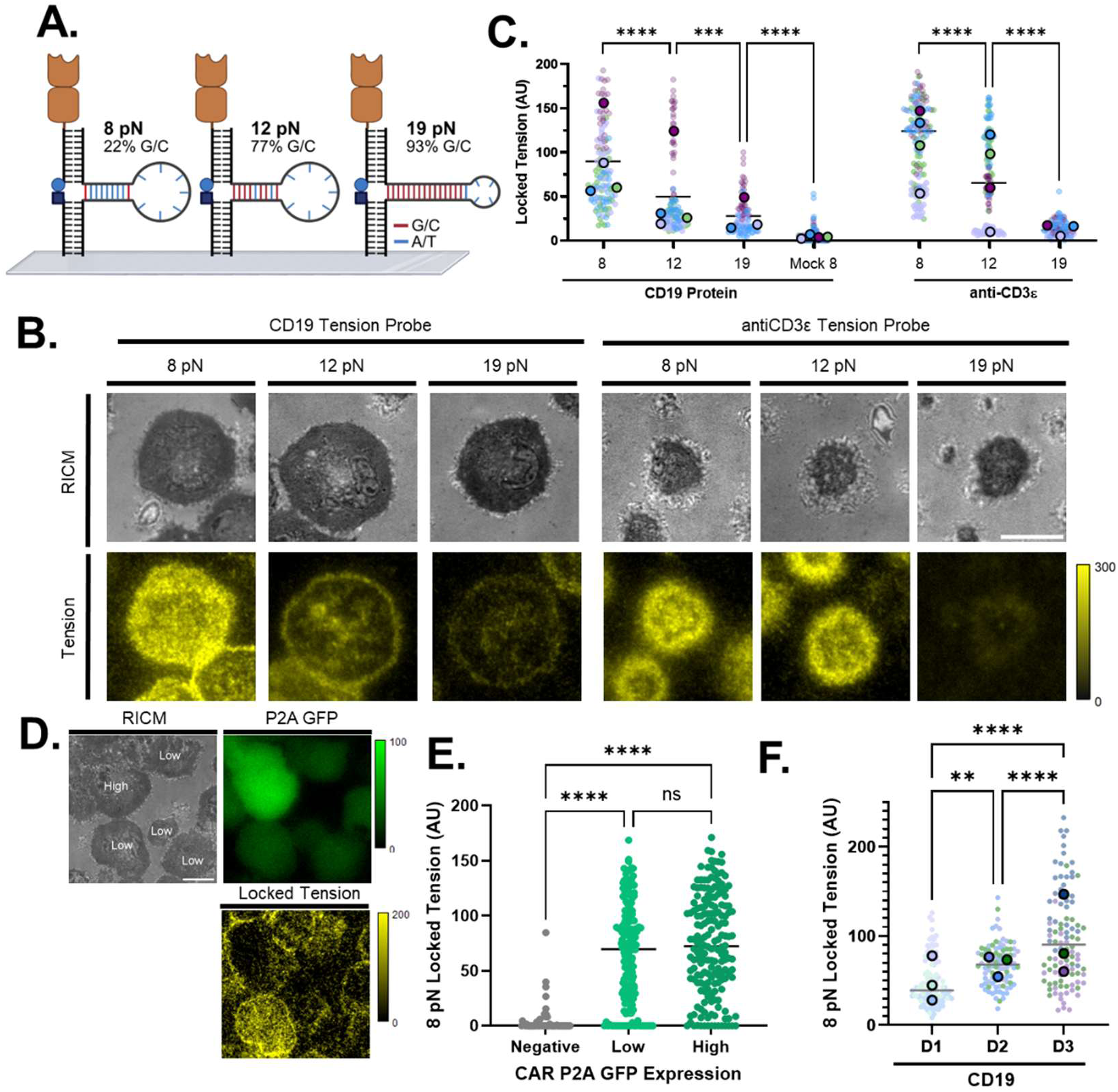
CAR T Cell Strength and Impact of Expression Level. **A**. Schematic showing increasing G/C content in stem region of hairpin which leads to greater values of F1/2. **B.** Representative RICM and CAR tension images showing decrease in locked 8 pN tension signal for greater F1/2 probes. **C**. Locked 8 pN tension signal for each F1/2 threshold. Data obtained from n=90 cells per condition from 3 transductions using the same human donor. Color encoded for each replicate and the larger data points show the mean of each transduction. **** <0.0001, *** = 0.0002. **D**. Representative RICM, 8 pN locked tension, and P2A GFP CAR channel showing the expression level of the CAR. The RICM image was labeled to indicate GFP signal for each cell. **E**. Plot showing 8 pN locked tension signal as a function of CAR expression level. n=60 cells (per bin) that were pooled from 2 transductions using 2 unique human donors. ****<0.0001. ns = 0.2086. **F**. Plot showing donor-to-donor differences in tension signal when CAR T cells were challenged using the same antigen, n > 90 cells/donor from 3 unique transductions per donor. **** <0.0001. ** = 0.0019.

We next aimed to determine if CAR copy number controls the CD19 tension signal measured in our assays. For example, if forces are primarily driven by CAR expression, cells that express more CAR will have more tension signal, but if the force generation by the cytoskeleton represents the limiting factor, then changes in CAR expression level would show minimal impact on the tension signal. Using a P2A GFP CAR to quantify CAR expression, we analyzed cells on the CD19 tension probe surface that were CAR^−^, CAR^low^ and CAR^high^ via their GFP expression (Figure 2D). Within these three populations, higher CAR expression did not cause changes in the amount of tension exerted, suggesting that within the average variability of a population, expression level is not the primary driver of higher tension measurements (Figure 2E). Other factors, such as actin recruitment might instead be the primary drivers of tension measurements.

Experiments in Figure 1 and 2 used CAR T cells proliferated from different transductions isolated from the same donor. Given the innate biological differences between healthy donors, we next wondered whether the CAR T tension signal could serve as a biomarker (mechanophenotype) for any given individual. To address this question, we measured CAR-CD19 tension from three individuals (D1, D2, D3) that provided healthy T cells at different points over the span of 6 months. We found that each transduction generated CAR T cells with differing mechanical activity, but individual donors showed distinctive mechanophenotypes, with 1.43 fold and 1.72 fold higher CAR-CD19 tension for D2 and D3 across transductions compared to D1 (Figure 2F). The data shows significant person-to-person variations in tension signal, and the reasons for this are currently unclear. We hypothesized that this may be due to difference in the underlying T cell activity, which motivated the subsequent investigations in how altered T cell health influences CAR T force transmission.

### Forces are Dependent on T Cell Fitness

We next examined the impact of T cell fitness state on CAR T cell forces (Figure 3A). Persistent exposure to antigen, immunosuppressive microenvironments, and tonic signaling from CAR T cells themselves all can induce T cell exhaustion, a state characterized by decreased proliferation and reduced cytotoxic capacity.^29–31^ To mimic exhausted CAR T cells *in vivo*, we followed an exhaustion model described in literature precedent and split a transduction into two groups, handling one group using conventional methods and continuously activating the other group with CD3/CD28 activation beads (Figure 3B). ^32^ This protocol allowed us to generate both PD-1^−^ healthy and PD-1^+^ exhausted CAR T cell populations from the same human donor source and the same transduction pool (Figure 3C, SI Figure 6).^32^ This is an important aspect of this model, as Figure 2 already showed that CD19-directed CAR forces are significantly dependent on both the donor and transduction.

**Figure 3.**
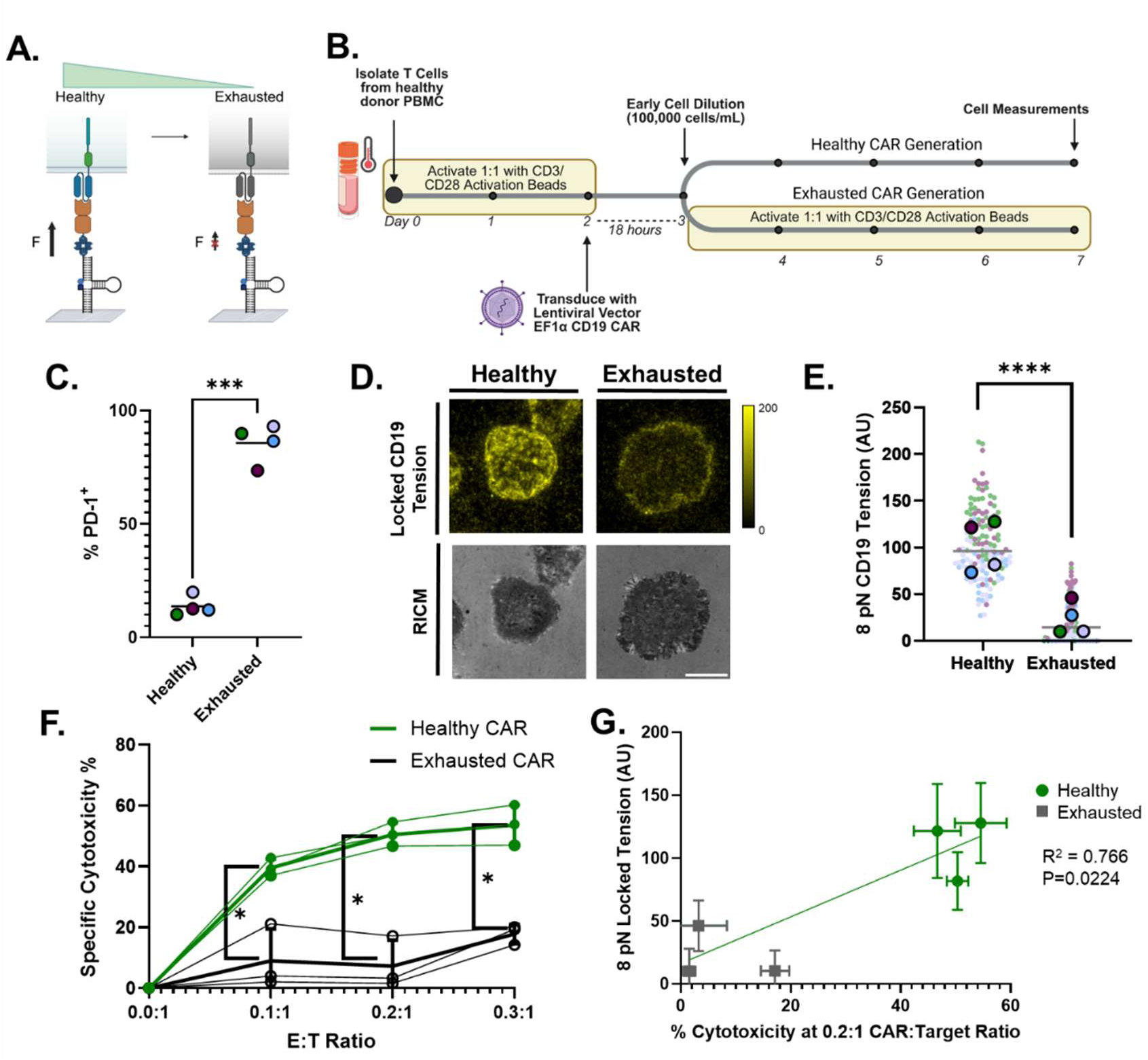
Varying T Cell Fitness Leads to Differential CAR Tension. **A**. Schematic of hypothesis that healthy CAR T cells generate greater forces compared to exhausted cells. **B.** Timeline of exhaustion model used in this work. **C**. Flow cytometry showing generation of a healthy PD-1^−^ CAR population and a PD-1^+^ exhausted CAR population. N=4, 10,000 cells/condition. P=0.0004, Student’s Two-Tailed Paired T Test. **D**. Representative RICM and 8 pN locked tension images showing healthy CAR T cells exert greater levels of tension compared to exhausted cells. **E**. Quantification of 8 pN locked tension for healthy and exhausted CAR T cells, 30 cells/donor, n=4 donors. P<0.0001, Student’s Two-Tailed Paired T Test. **F.** Healthy and exhausted cell specific cytotoxicity, n=3 independent cytotoxicity assays from 2 unique donors. P= 0.0219, 0.0170 .0222, Student’s Two-Tailed Paired T Test. **G**. Plot showing strong relationship between cytotoxicity and tension measurements. R^2^ = 0.6925, P = 0.0399.

We measured CD19-CAR forces in these healthy and exhausted populations and found that exhausted T cells showed a 78.9% decrease in 8 pN forces compared to healthy CARs (Figure 3D and 3E). Since exhausted T cells are known to have decreased effector functions, we next sought to establish an association of CAR T cell forces with cytotoxicity. To accomplish this, we co-cultured CD19 CAR T effector (E) cells with RS4;11 target (T) cells, a CD19-positive B-ALL cell line, at varying E:T ratios. The target cells were pre-labeled with VPD450 to provide a bright and long-lasting target cell label for flow cytometry. Cells were stained with both eFluor780, an amine reactive viability dye, and Annexin-V APC, a reporter of early apoptosis via phosphatidyl serine, and cytotoxicity was measured by positivity for either marker on target cells (SI Figure 7). The CAR-specific cytotoxicity was assessed by subtracting the background cell killing of the corresponding mock condition from both healthy and exhausted CAR T cells. Exhausted CAR T cells had decreased CAR-specific killing activity compared to healthy CARs across all E:T ratios (p<0.05 for all cell groups) (Figure 3F). This decreased killing matched decreased tension measurements (Figure 3G), showing a strong association between tension and cytotoxicity.

### CAR T Cell Forces are F-Actin-Mediated and Independent of Nck

TCR-antigen forces are mediated by actin dynamics, and inhibiting actin activity has been shown to modulate these forces. To investigate the source of CAR forces, we selected small molecule inhibitors shown to dampen TCR forces by impacting the cell’s actin recruitment and polymerization (Figure 4A). We seeded CAR T cells on 8 pN DNA hairpin tension probes displaying CD19 for 10 minutes to allow for efficient spreading, then treated them with actin inhibitors to change their real time tension before adding locking strand (Figure 4B). As expected, Latrunculin B, a potent inhibitor of F-actin, and CK666, an inhibitor of Arp2/3, completely abolished tension, showing that CD19 forces are actively mediated by recruitment and remodeling of the actin cytoskeleton (Figure 4C).

**Figure 4.**
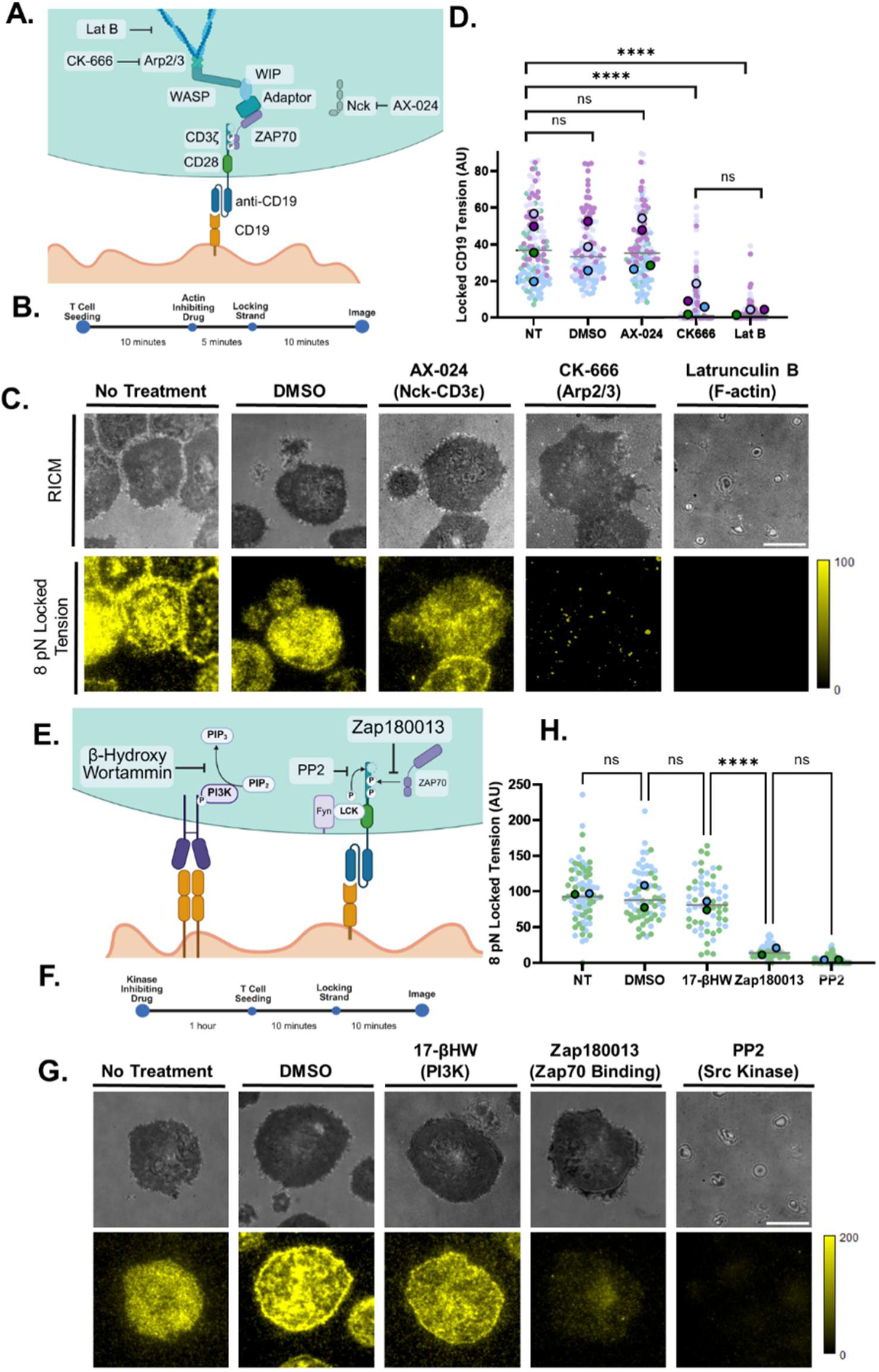
Effects of actin inhibitors and kinase inhibitors on CAR T cell tension generation. **A**. Schematic visualizing how a CAR could couple to the actin cytoskeleton and mechanisms of inhibitors. **B**. Timeline of drug conditioning experiments for actin inhibitors. **C**. Representative RICM and 8 pN locked tension images (Cy3B) showing the effect of actin inhibitors on CAR T cell spreading and tension. **D.** Plot quantifying single cell tension for different treatment groups (500 nM AX-024, 50 μM CK666, 5 μM LatB). n>90 cells/condition across 3-4 transductions. ****<0.0001. **E**. Schematic postulating how kinase inhibitors modulate CAR cytoskeletal coupling. **F**. Timeline of drug conditioning experiments for kinase inhibitors. **G.** Representative RICM and 8 pN locked tension images (Cy3B) showing the effect of actin inhibitors on CAR T cell spreading and tension. **H**. Plot quantifying single cell tension for different treatment groups (50 nM PP2, 10 μM Zap180013, or 50 nM 17β-Hydroxywortmannin). n>60 cells/condition across 2 transductions. ****<0.0001.

Interestingly, Nck inhibition causes a 42.5% reduction of aCD3ε tension for the native TCR (SI Figure 8), but did not alter forces exerted by the CAR (Figure 4D), showing that a CAR T cell’s cytoskeletal coupling is Nck-independent. Since CARs lack the CD3ε domain, and Nck is shown to interact with this domain, it is not surprising that inhibiting this interaction failed to show significant changes in CAR forces.^33^ This further shows the differences between forces generated by CAR T cells and those generated by native T cells.^34, 35^

Given that TCR activation is required for force generation, we next sought to study how proximal kinase activity mediates CAR forces (Figure 4E). We pretreated CAR T cells using small molecule kinase inhibitors for 1 hour, and after seeding cells on the surface and allowing for cells to spread, locking strand was added for 10 minutes and tension was measured (Figure 4F). Interestingly, 17-βHW inhibition of PI3K, which can be recruited to the intracellular CD28 domain of CAR T cells, had no detectable effect on tension (Figure 4G, 4H). While it was previously reported that the full CD28 protein is mechanically sensitive, the intracellular domain utilized for CAR T cells is not sufficient to create the mechanical response of the CAR.^18^ Conversely, inhibition of CD3ζ phosphorylation via the receptor tyrosine kinase (RTK) inhibitor PP2 caused strong inhibition of tension measurements (Figure 4G, 4H), showing that phosphorylation of the ζ chain is necessary for tension generation and strengthening the link between CAR activation and forces.

### Modulation of Tension with Small Molecule Inhibitors and Altered CAR Constructs

Based on the strong relationship between kinase inhibition and force generation, we next looked at a RTK inhibitor known to have a potent effect on CAR T cell cytotoxicity. Dasatinib, a broad spectrum SRC kinase inhibitor, has inhibitory effects on Src, Fyn and Lck, signaling kinases that are important for early T cell activation and function, and has been used clinically to inhibit BCR-ABL fusion protein in chronic myeloid leukemia (CML).^36–38^ Recent literature has shown that the conserved signaling pathways between TCRs and CARs allow dasatinib to potently suppress CAR T cell activation and cytotoxicity (Figure 5A).^39, 40^ Thus, we investigated the impact of dasatinib on CAR T force transmission.

**Figure 5.**
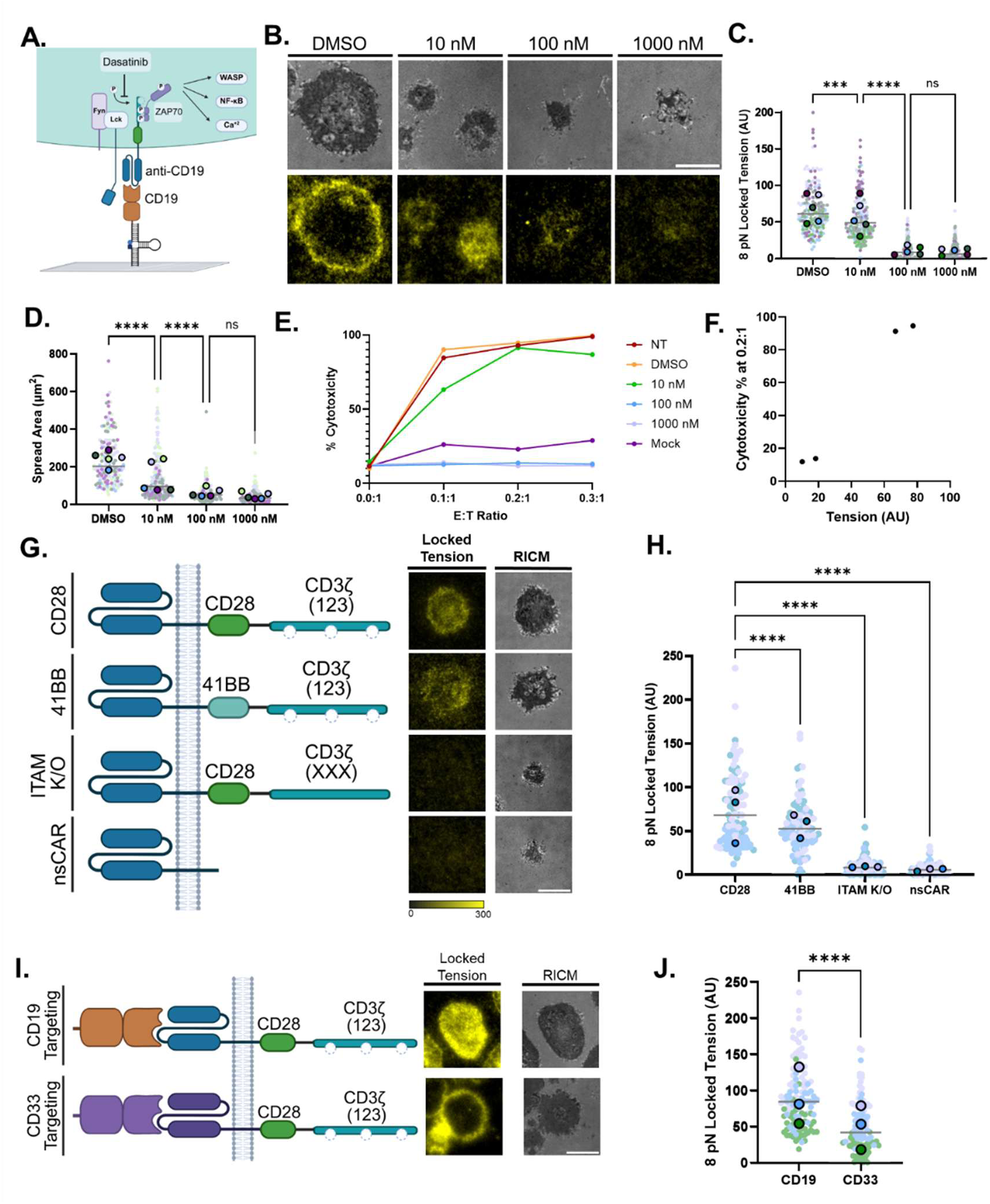
Cytoskeletal coupling to CAR cytoplasmic domain is phosphorylation-dependent. **A.** Proposed mechanism of action for dasatinib in CAR T cells which leads to Lck inhibition. **B**. Representative RICM and tension images of CAR T cells treated with escalating doses of dasatinib. **C** and **D.** Plots quantifying CAR T cell forces and cell spread area as a function of dasatinib treatment. N ≥ 162 from 5 independent transductions. ***= 0.0001, ****<0.0001. **E**. Plot showing cytotoxicity as a function of dose-dependent dasatinib CAR T cell treatment. Data is averaged from two technical replicates using the same batch of CAR T cells. **F**. Relationship between CAR T cell tension and cytotoxicity showing correlation. **G**. Schematic showing three additional engineered CARs with 4-1BB costimulatory domain, ITAM mutants lacking key tyrosine residues, and finally CAR lacking a cytoplasmic domain along with representative RICM and 8 pN locked tension (Cy3B) images. **H**. Plot of single cell tension levels for each CAR T cell construct tested. N ≥ 162 from 5 independent transductions. One-way ANOVA. ***= 0.0001 , ****<0.0001. **I**. Representative RICM and 8 pN tension images showing the tension and spread area of cells on CD19 and CD33 antigens. **J.** Quantification of cellular tension levels from n≥90 cells/condition from 3 separate transductions. Each color indicates a biological replicate from a unique transduction using the same human donor. The larger data points indicate the mean of each replicate while the small data points represent the single cell measurements. N ≥ 106 cells/condition from 3 independent transductions. Student’s Two-Tailed Paired T Test, ****<0.0001.

24 hours before functional testing, we dosed T cells with either vehicle or dasatinib at 10, 100 or 1000 nM as described previously, and conserved these conditions for tension and cytotoxicity assays.^39^ Increasing doses of dasatinib led to decreases in tension and spread area in a dose-dependent manner across transductions with different donors (Figure 5B). At 10 nM dasatinib, a 19.4% reduction in CAR tension signal was observed, which was increased to a 76.5% reduction in tension at 100 nM and an 84.8% reduction at 1000 nM (Figure 5C). These results validate the role of proximal kinase in mediating CAR mechanotransmission observed in Figure 4. Similar trends were noted in the spread area of cells, with higher doses of drug corresponding to decreased spread area (Figure 5D). We next conducted cytotoxicity assays following the protocols used to obtain data in Figure 3F and assessed changes in cytotoxicity across E:T ratios and drug dosages. We found that high doses of dasatinib efficiently shut down CAR T cell cytotoxicity at all E:T ratios, with CAR T cells treated with 100 or 1000 nM dasatinib having similar target RS4;11 cell death as that measured for conditions lacking CAR T cells (Figure 5E). In agreement with literature, this multikinase inhibition using dasatinib abolished CAR T killing to undetectable levels (Figure 5E). When plotting CAR-specific forces against cytotoxicity measurements, we were again able to see a strong relationship between tension and cytotoxicity (Figure 5F).

We next investigated the coupling between the cytoskeleton and the CAR by genetically modifying altered cytoplasmic CAR domains. 2^nd^ generation and later CAR T cells contain a costimulatory domain that signals for increased T cell proliferation, cytotoxic function, memory formation or survival.^41, 42^ Although several co-stimulatory domains have been used, the CD28 and 4-1BB coreceptors are most commonly used clinically.^43^ The intracellular domains have distinct functions: CD28 recruits PI3K and Lck and is linked to increased cytokine production, whereas 4-1BB recruits TRAF proteins and is associated with prolonged *in vivo* persistence.^43^ ITAM knockouts containing tyrosine to phenylalanine mutations in the 3 ITAMs of the CD3ζ chain of the CAR have also been used to prevent CAR T cell signaling.^44^ To study CAR T cell binding without inducing cytotoxicity, CARs lacking internal domains to signal have also been developed.

Accordingly, we also designed a CD19-41BB-CD3ζ CAR (41BB), a CD19-CD28-CD3ζXXX CAR (ITAM K/O) and a CD19 non-signaling CAR (nsCAR) to investigate the roles of co-stimulation, ITAMs, and the entire cytoplasmic signaling domain on tension (Figure 5G). Interestingly, the 41BB CAR showed a decrease in average mechanotransmission compared to the CD28 CAR, though both were able to transduce tension (Figure 5H). In support of this finding, previous work has found CD28 costimulatory domains can directly recruit Fyn kinase, whereas 4-1BB is unable to, driving weaker CD3ζ phosphorylation.^45^ These results are consistent with both our findings that 4-1BB drives weaker tension and that Src family kinases (as inhibited by PP2 and dasatinib) are important for CAR T cell tension. The ITAM K/O CAR and nsCAR, which cannot become phosphorylated, had a striking 87.0% and 91.5% decrease in tension signal respectively (Figure 5H), further underscoring the necessity of ITAM phosphorylation to generate tension, which is also consistent with our observations upon kinase inhibitor treatment.

Finally, we were interested in seeing if CAR-CD19 forces were unique to CD19 or if the transmission of biophysical forces was a general feature of CAR-antigen complexes. We chose to study a CD33 scFv that is part of a CD33-CAR under investigation (phase I clinical trial: NCT03971799) as a therapy for acute myeloid leukemia (AML).^46^ The scFv is derived from the anti-CD33 humanized monoclonal antibody HuM195.^47^ The CD33-CAR was chosen due to its similar affinity for antigen (K_d_ = 4.75 nM) as the CD19-directed CAR’s affinity to CD19 (2-6 nM), and was designed with an identical promoter, costimulatory and signaling domains.^48, 49^ Tension measurements using the 8 pN lock probe showed clear tension patterns for CD33-CARs (Figure 5I) with timescales of accumulation and spatial patterns that were consistent with that of CD19-CARs (SI Video 3). However, despite the structural parallels between the two CARs, the magnitude of tension generated by the CD33-CAR was 45.6% lower than that of the CD19-CARs when tested using the same donor and T cells (Figure 5J). This suggests that these CAR-antigen forces are conserved, but the magnitudes of accumulated tension are antigen specific. Importantly, given that CD33 and CD19 affinity for target is nearly identical, this indicates that mechanical measurements reflect other parameters such as bond lifetime under force, signaling and cytoskeletal dynamics.

## Discussion

This work reveals and quantifies the pN forces transmitted between CARs and their antigens. This was a previously underappreciated aspect of CAR T cell signaling and function, and shows these forces are present when CAR T cells are functional with the frequency of mechanical events tightly linked to their cytotoxicity. Utilizing DNA hairpin tension sensors, we were able to visualize specific and robust forces for CD19 CAR with brief, 1s mechanical events that ranged in magnitude up to 19 pN. CAR forces showed donor-to-donor variations, with some donors having a stronger “mechanophenotype” across several transductions, suggesting that the ability to generate mechanical force could be reflective of the functional potency of specific T cell populations. We show these forces are highly sensitive to the functional state of the T cell, with exhausted T cells having abrogated forces and cytotoxic function. We further found that both conventional actin inhibitors and kinase inhibitors dampened CAR T cell forces, suggesting the necessity of actin polymerization, Arp2/3 and proximal kinases such as Zap70, Lck, and Fyn to generate force. We utilized a dose-dependent kinase inhibitor, dasatinib and observed a strong link between force and cytotoxic effect in CAR T cells. Finally, by engineering individual domains of the CAR, we discovered that functional ITAMs are required for force generation and noted the role of the CD28 coreceptor in increasing force transmission, pointing to a coupling between CAR T cell activation and force generation that requires phosphorylation of ITAMs to recruit actin adaptors for mechanical coupling. Interestingly, CAR-antigen forces are more transient and less frequent compared to that of TCR-pMHC events, which is clear when comparing real time signals.^11^ This is likely why CAR forces were not previously reported despite long-standing speculation and underscores the importance of the locking strand to increase signal-to-noise. Because the CAR represents a minimal fragment of the TCR complex, it allows for identifying functional elements of the TCR that mediate force transmission. Specifically, our work suggests that the pY ITAM domains of CD3ζ represent critical sites of cytoskeletal coupling as the phosphorylation of these sites is required for mechanotransmission. A limitation of this study is the use of the immobilized probes to create maps of CAR T cell forces, which might localize differently at a true cell-cell interface due to the CAR T cell’s ability to laterally translocate CD19 antigen. Nevertheless, these CAR T cell forces are well correlated with the functionality of CAR T cells and the use of tension probes establishes a rapid method for testing the mechanical strength and functionality of CAR T cells. Beyond the fundamental significance of revealing biophysical aspects of CAR signaling, we anticipate potential applications in predicting CAR T cell activity when screening new CAR-antigen therapies and when determining the potency of CAR treatments in a personalized manner.

## Materials and Methods

### 1. Materials

#### 1.1. Oligonucleotides

All oligonucleotides were custom synthesized by Integrated DNA Technologies (Coralville, IA). Table S1 includes the names and sequences of all strands used in this work.

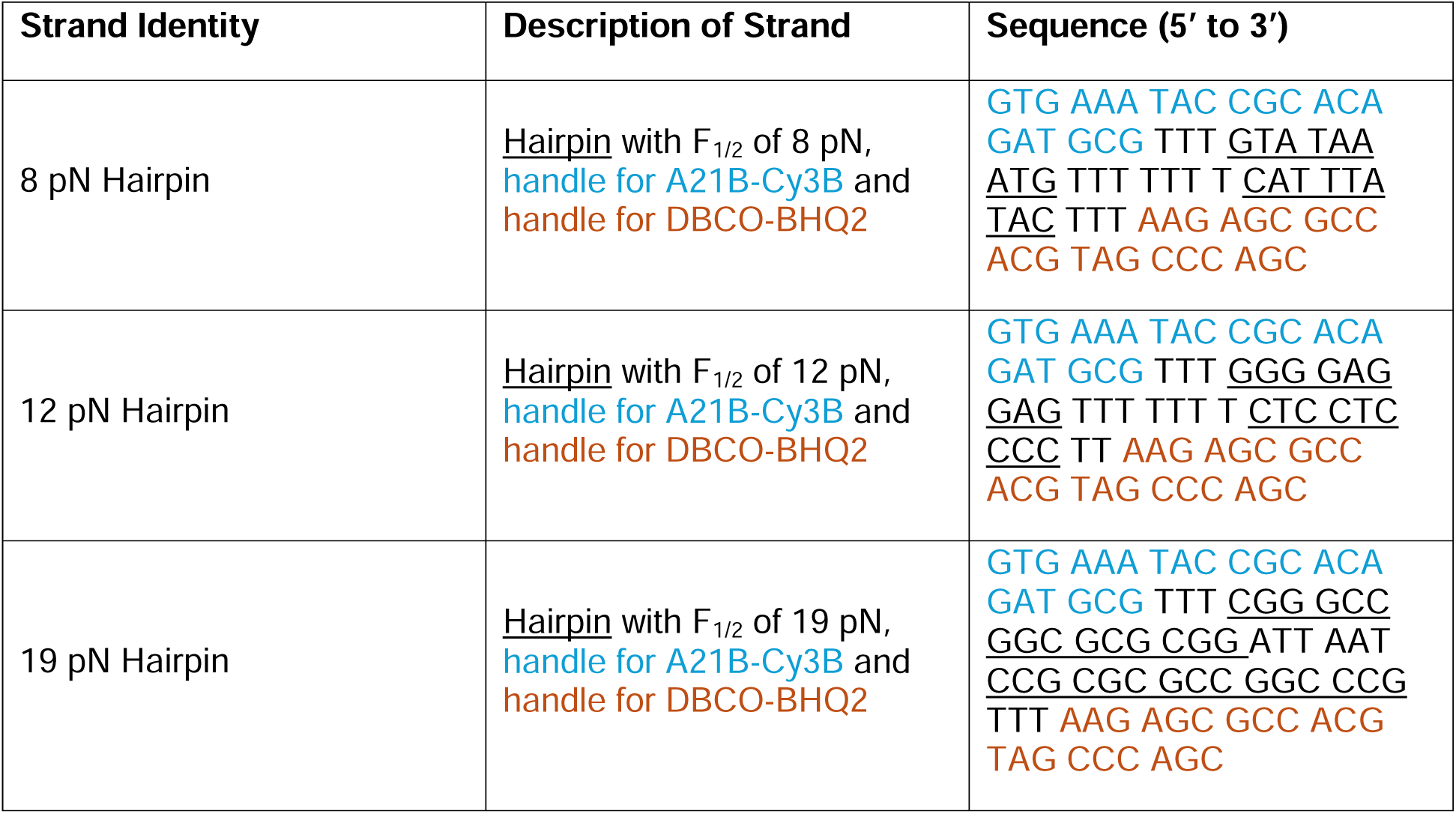
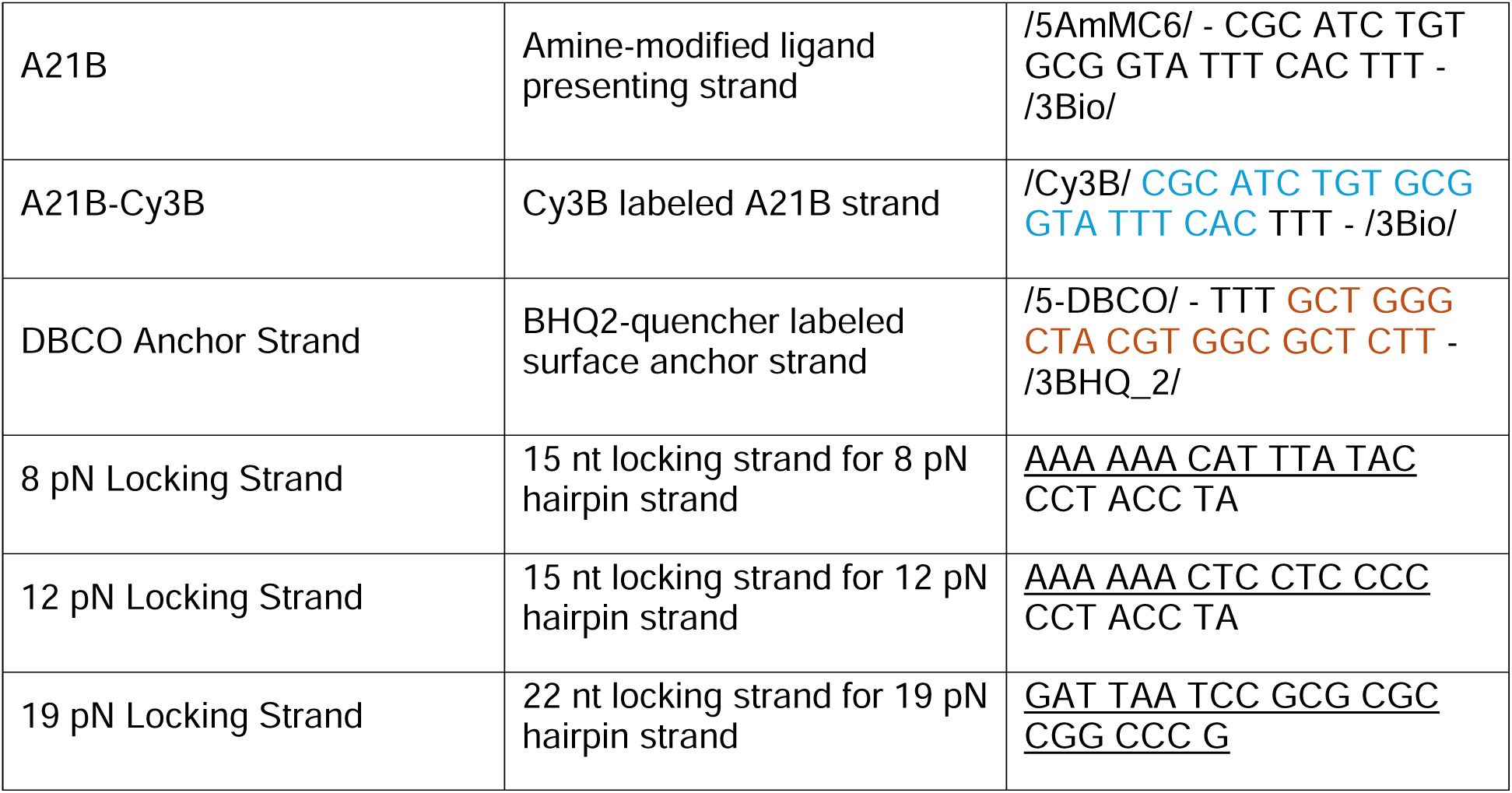

#### 1.2. Consumable Reagents and Antibodies

3-Aminopropyltriethoxysilane was purchased from Acros (Cat# AC430941000, Pittsburgh, PA). NHS-PEG4-Azide was purchased from BroadPharm (Cat#BP-20518, San Diego, CA). Sulfo-NHS acetate (Cat# 26777), Teflon racks (Cat# C14784), Biotinylated OKT3 CD3 mAb (Cat# 13-0037-82), 1 M MgCl_2_ (Cat#AM9530G), Human T Cell CD3/CD28 activation Dynabeads (Cat# 11161D) and eBioscience Fixable Viability Dye eFluor 780 (Cat#501129035) were purchased from Thermo Fisher Scientific (Waltham, MA). Streptavidin (Cat# S000-01) was purchased from Rockland Immunochemicals Inc. (Rockland, NY)Ethanol (Cat# 459836), hydrogen peroxide (Cat# H1009), bovine serum albumin (BSA) (Cat# 10735078001), and Hank’s balanced salts (H8264) were purchased from Sigma Aldrich (St. Louis, MO). ProPlate Bottomless Microtiter Plates (Cat#204969) were purchased from Grace BioLabs. Cy3B NHS ester (Cat# PA63101) was purchased from GE Healthcare (Pittsburgh, PA). Biotinylated CD19 protein (Cat#CD9-H82E9-25ug), biotinylated CD33 protein (Cat#CD3-H82E7-25ug), and AlexaFluor 647 Anti-FMC63 Ab (Cat#FM3-AM534-25tests) were purchased from AcroBioSystems (Newark, DE). FITC anti-human CD279 (PD-1) Antibody (Cat# 329903), and APC Annexin V (Cat#640941) were purchased from BioLegend (San Diego, CA). Violet Proliferation Dye 450 (Cat# 562158) was purchased from BD Biosciences (San Jose, CA). Ficoll-Plaque PLUS density gradient (Cat# GE17-1440-02) was purchased from GE HealthCare (Chicago, IL). Polybrene (Cat# TR-1003) was purchased from EMD Millipore (Billerica, MA). Human IL-2 (Cat#AF-200-02-250UG_61262816) and human IL-7 (Cat# AF-200-07-10UG_61263190) were purchased from PeproTech (Carlsbad, CA). X-Vivo 15 (Cat#04-418Q) was purchased from Lonza (Basel, Switzerland). Dasatinib (Cat #73082) and EasySep Human T Cell isolation kit (Cat# 17951) were purchased from StemCell Technologies (Vancouver, BC). Latrunculin B (L5288-1MG) was purchased from Millipore Sigma (Burlington, MA). CK666 (Cat#ab141231) was purchased from Abcam (Cambridge, United Kingdom). AX-024 (Cat# S6727) was purchased from selleckchem (Houston, TX). PP2 (Cat# HY-13805) and Zap180013 (Cat#HY-136179) were purchased from MedChemExpress (Monmouth Junction, NJ). 17β-hydroxy Wortmannin (Cat# sc-358740) was purchased from Santa Cruz Biotech (Santa Cruz, CA).

Custom lentiviral packaging was done by VectorBuilder (Chicago, IL). All buffers were made with Nanopure water (18.2□MΩ).

#### 1.3. Cell Lines

CD19 expressing RS4;11 cell line was gifted to us by the laboratory of Dr. Douglas Graham (Emory University, Atlanta, GA, USA). Cells were maintained at a concentration of 0.5×10^6^–2×10^6^ cells/mL in RPMI-1640 with L-glutamine (Corning CellGro, Manassas, VA, USA) supplemented with 10% fetal bovine serum (FBS; R&D Systems, Minneapolis, MN, USA) and 1% Penicillin-Streptomycin (Thermo Fisher Scientific, Waltham, MA, USA).

#### 1.4. Human T Cell Isolation

Peripheral blood from healthy adult volunteers was obtained under an IRB approved protocol (IRB00101797) through the Emory University Children’s Clinical and Translational Discovery Core (CCTDC), after which peripheral blood mononuclear cells (PBMCs) were isolated using a Ficoll-Plaque PLUS density gradient and cryopreserved. T cells were negatively selected from thawed PBMCs using the EasySep^TM^ Human T cell Isolation Kit (STEMCELL Technologies, Vancouver, Canada).

### 2. Equipment

#### 2.1. Microscopes

The microscope used for imaging was a Nikon Ti-2 Eclipse inverted epifluorescence microscope with an Andor 512 EMCCD Camera, a Lumen Dynamics X-Cite 120 LED light source, a CFI Apo 100× NA 1.49 objective (Nikon), and a TIRF launcher with four laser lines: 405 nm (20 mW), 488 nm (80 mW), 561 nm (80 mW), and 647 nm (125 mW). All of the reported experiments were performed using the following Chroma filter cubes: GFP, TRITC, Cy5, and RICM.

#### 2.2. Flow Cytometers

The spectral flow cytometry for cytotoxicity assays was completed on 4- (V B Yg R) or 5- (UV V B Yg R) laser Cytek Auroras in the Emory Pediatrics/Winship Flow Cytomtery core. Spectral Unmixing (deconvolution) was completed in the SpectroFlo software on unstained and singly stained controls.

Exhausted cell profiling was completed on a Beckman Coulter Cytoflex (V0-B2-R2) with 488 nm and 638 nm lasers.

### 3. Methods

#### 3.1 CD19-CAR T Cell Generation

The complementary DNA (cDNA) sequence for the FMC63-based CD19-CAR was codon optimized for optimal expression in human cells using the codon optimization tool from IDT (Coralville, IA). The codon-optimized gene block for the CD19-CAR cDNA sequence was created by Genewiz (South Plainfield, NJ) and was subsequently cloned into a vector backbone with an EF1α promoter given to us courtesy of the laboratory of Dr. H Trent Spencer (Emory University, Atlanta, GA, USA). The CAR construct was a second-generation CAR with a CD28 co-stimulatory domain as depicted in Figure S1. The CAR-encoding plasmid was sent to VectorBuilder (Chicago, Illinois, USA) for high-titer 3^rd^ generation lentiviral vector production using a four-plasmid system. Titer for vector produced ranged between 10^8^-10^9^ TU/mL.

Isolated human T cells from healthy donors as described in section 1.4 were stimulated with anti-CD3/CD28 Human T-Activator Dynabeads at a 1:1 bead-to-cell ratio in X-VIVO supplemented with 10% FBS, 1% penicillin-streptomycin, 0.1 µL/mL human IL-2 and 0.1 µL/mL human IL-7 at ∼2 × 10^6^ cells/mL. After 48 hours of stimulation, beads were removed. Approximately 1 × 10^6^ stimulated T cells were transduced with len iviral vector at multiplicity of infection (MOI) 20 with 10 μg/mL polybrene for 18 hours. MOI was determined using the following calculation:

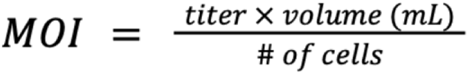

After 18 hours, cells were resuspended in fresh media and diluted to ∼1 × 10^5^ cells/mL, then left to expand for 96 hours before CAR expression was determined and cells were used in downstream studies.

#### 3.2. Labeling of Oligonucleotides

A mixture of A21B (10 nmol) and excess Cy3B NHS ester in 0.1 M sodium bicarbonate and 10% DMSO was allowed to react at room temperature for 1 hour. The reaction was purified with P2 gel filtration to remove unreacted dye, and further purified to a solely-Cy3B modified product by reverse-phase HPLC in an AdvanceBio Oligonucleotide column (solvent A: 0.1 M TEAA; solvent B: 100% ACN); the initial condition was 10% B with a gradient of 1%/min and a flow rate of 1 mL/min.

#### 3.3. Surface Preparation

No. 2 glass coverslips were rinsed and sonicated with nanopure water (18.2 MΩ cm^−1^) for 20 min, and then sonicated with 200 proof ethanol for 20 min and rinsed 6 times with nanopure water to remove ethanol. The cleaned slides were submerged in fresh piranha solution (3:1 v/v = H_2_SO_4_: H_2_O_2_) used to clean the substrates for 30 min. **Caution: Pirahna is extremely reactive and explosive if in contact with organic solvents.** Afterwards, the substrates were rinsed 6 times with nanopure water. The substrates were then sonicated in 200 proof ethanol to remove excess water. The slides were next modified to display amines with a 3% v/v APTES solution in ethanol for 45 min. The amine-modified coverslips were then rinsed in ethanol 3 times and dried under a stream of N_2_ gas and placed in an oven for 20 min at 80°C. The surfaces were then adhered to ProPlate Bottomless Microtiter plates (Grace) and reacted with NHS-PEG_4_-Azide in 0.1 M sodium bicarbonate for 1 hr to install an azide functional group on the glass. The slides were then blocked with sulfo-NHS acetate in 0.1 M sodium bicarbonate for 30 min to block remaining amine sites. The DNA tension probe hairpins were assembled in 1x PBS + 1 M NaCl by mixing the Cy3B labeled A21B strand (400 nM), hairpin strand (400 nM) and anchor strand (440 nM) in the ratio of 1 : 1 : 1.1. The solution was then heated to 75°C for 5 min by using a thermocycler and cooled back to room-temperature over a period of 20 min. 50 µl of this final solution was added to each well for a final concentration in the well of 200 nM tension probe, which was incubated overnight at 4 °C. DNA tension probe modified coverslips were rinsed in 1x PBS 3 times before further functionalization with 40 µg/ml of streptavidin in 1X PBS for 1 hr. The coverslips were rinsed in 1x PBS to remove the unbound streptavidin, which was followed by the final modification with 20 µg/ml of biotinylated CD19 or CD33 or 40 µg/mL antibody in 1X PBS for 1 h. The coverslips were rinsed in 1x PBS to remove excess ligand, buffer exchanged to HBSS and used immediately for experiments.

#### 3.4. Fluorescence Microscopy

The microscope used was NSTORM Nikon microscope (Ti E motorized inverted microscope body), operated by Nikon Elements software. A CFI Apo 100X NA 1.49 objective was used and a 512 iXon Ultra EMCCD camera (Andor). The optical system includes a total internal reflectance fluorescence (TIRF) variable mirror launcher and a Nikon Perfect Focus System, an interferometry-based device which corrects z-drift of the stage. Reflection interference contrast microscopy (RICM) and epifluorescence images were captured a Lumen Dynamics X-Cite 120 LED light source. All of the reported experiments were performed using the following Chroma filter cubes: RICM (97270 SRIC C168785), GFP, TRITC, and Cy5.

Imaging was performed using Hank’s Balanced Salts supplemented with 8 mM MgCl_2_. All imaging data was acquired at room temperature. Images in TRITC were taken with 500 ms exposure time with no EMCCD gain.

#### 3.5. Real Time Single Molecule Imaging and Analysis

Images were taken with 500 ms exposure time at ∼0.75 Hz over 5 minutes, with a representative fluorescent image of frame #50 shown in **Figure S2A**. To deconvolute real-time tension events from a steady-state fluorescent background, a background subtraction was performed by first subtracting a moving average of 10 frames using a subset of scripts from SMALLLABS software^1^, which sets the average fluorescent value to 1000 AU (**Figure S2B**). Afterwards, a subsequent rolling ball subtraction with a radius of 50 pixels using Fiji was performed to aid downstream puncta localization (**Figure S2C**). Once background subtracted, the fluorescent puncta were localized using ThunderSTORM software (**Figure S2D**).^2^ Sequential events within 350 nm were treated as a single, long-lived tension event when obtaining the lifetime of tension events,^3^ while all other localizations were assigned a tension lifetime of one frame. To approximate the photobleaching rate of an opened probe under our optical conditions a surface labeled with a sparse density Cy3B was prepared (**Figure S2G**). Fluorescence signal of this surface was measured using identical optical settings, and the average background intensity was fit to an exponential decay + constant baseline to obtain an approximate photobleaching rate of 138.7 s (105 frames) (**Figure S2H**). Finally, the tension lifetimes were binned, and the probability distribution function (PDF) was fit to an equation of the form: 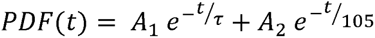 (**Figure 1F - Manuscript**).

The single-molecule nature of detected events can be validated by both 1) the Gaussian shape of the fluorescent puncta and 2) a digital on/off nature of the fluorescent signal. We show a validation of the Gaussian shape of each puncta in SI Video 2, where we show the RICM and background-subtracted fluorescent signal of the same ROI as shown in Figure 1D (frame 50). In the video, we randomly selected 8 detected puncta from the ROI and marked them with a 5×5 blue square before zooming into the region and rotating the field of view to allow visualization of their Gaussian shape (SI Video 2). To validate the on-off nature of the fluorescent signal we analyzed the fluorescent traces and formed a pool of all tension events lasting exactly 3 frames (N = 336 events). From those events we randomly selected 10 traces and plotted their background-subtracted fluorescent signal normalized to a y-axis of 0 to 1 (**Figure S2E**). As these are single-molecule events they should be analyzed on an individual scale, but an averaging of these traces can aid visualization of the on-off fluorescent signal for these events by reducing spurious noise (**Figure S2F**).

We also sought to mitigate the effect of spurious “overlapping events” which would result in artificially high observed tension lifetimes. For any live-cell time trace with a given density of events, there is some probability that uncorrelated events in subsequent frames would be interpreted as the same event, simply because they occurred within close spatial-proximity to one another. One can imagine recording a tension event at t = 0, waiting a functionally infinite time, and then recording a tension event at the same location at t = ∞, leading to the obviously false conclusion that this tension event lasted for infinite time. To mitigate these spurious events in a controlled manner we downsampled our data to timelapses 20 frames apart (functionally infinite time compared to the CAR-CD19 tension lifetime). Analyzing these downsampled time traces yields an approximate distribution of observing subsequent tension events that are uncorrelated. Thus, when fitting the lifetime of tension events (Figure 1E) this distribution of falsely-correlated events was first subtracted from the measured tension lifetimes before fitting, minimizing their effect on the measurement.

Lastly, we would note that we actively compensate for false-positives in long lifetimes (such as the spurious overlapping events, as described above) but do not compensate for false-positives in short lifetimes (such as false signals which exist for 1 frame, missing puncta which cut longer events into several shorter events, or longer events which “turn off” between observations). Accounting for these effects quantitatively is much harder. This means that we likely observe more events that last for 1 frame than the underlying true value, and thus we describe our observed tension lifetime as a <u>lower bound</u> of the tension lifetime.

#### 3.6. Tension measurements with locking strand

Cells were allowed to spread on tension-probe covered surfaces for 10 minutes. After spreading, initial images were taken of real-time tension before the addition of 100 uL of 400 nM locking strand into 100 uL to a final concentration of 200 nM. Tension was allowed to accumulate for 10 minutes, and final images were taken in TRITC and RICM.

#### 3.7. Microscopy Data Analysis

Data analysis was completed in ImageJ on at least 30 cells per condition. Background subtraction was performed by selecting 3 regions of interest (ROIs) lacking cells in RICM channel, averaging their mean fluorescence values in Cy3B tension channel and subtracting this average background from the Cy3B tension channel.

#### 3.8. Exhausted CD19-CAR T Cell Generation

The exhausted CAR T cell phenotype was generated utilizing a previously established protocol.^4^ Briefly, healthy human T cells were isolated and transduced with CAR lentiviral vector as discussed in Sections 1.4 and 3.1. After the transduction, 10^6^ cells were separated out, diluted to 10^5^ cells/mL in X-Vivo, and incubated with CD3/CD28 activation beads in a 1:1 ratio for 3 days to induce exhaustion. The remaining healthy cells were rested in X-Vivo with cytokines for 3 days to create a healthy CAR population. Exhaustion was confirmed by surface PD-1 assessment by flow cytometry.

#### 3.9. *In vitro* cytotoxicity assay for healthy and exhausted CAR T cells

A flow-based cytotoxicity assay was utilized to measure the cytotoxicity of CAR T cells. Briefly, the healthy or exhausted mock (non-transduced) T cells or CAR T cells were co-cultured with 100,000 VPD-450 pre-stained CD19^+^ RS4;11s at effector to target (E:T) ratios of 0.1:1, 0.2:1 and 0.3:1 in a U bottom 96 well plate for 24 hours at 37°C with 5% CO2. The cells were stained with eFluor780 and APC Annexin V to determine live and dead cell populations. Samples were washed with 150 uL PBS, spun down at 320 × *g* for 5 min, then incubated with Annexin V and eFluor780 in Annexin V binding buffer for 30 minutes on ice. Cells that were VPD450^+^ eFluor780^−^ and APC Annexin V^−^ (SI Figure 5) were enumerated as the live cell percentage. At least 15,000 events were analyzed per condition in technical duplicates.

CAR-specific lysis was calculated by:

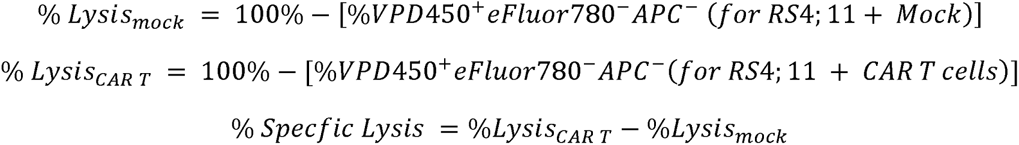

#### 3.10. PD-1 Staining and Flow Cytometry

Healthy and exhausted CARs were stained for 30 minutes on ice in 1:100 anti-PD-1 antibody in FACS buffer, followed by centrifugation (300 × *g*, 5 min) and resuspension in fresh FACS buffer prior to flow cytometry. At least 10,000 events were analyzed per condition.

#### 3.11. Actin Inhibitors

Actin inhibition was performed as previously described. Briefly, cells were allowed to spread and engage with the surface for 10 minutes to enable efficient cell spreading. Cells in 100 μL of HBSS were then drugged with either 1 μL of untreated DMSO or 1 μL of a 100x drug stock to a final concentration of either 500 nM AX-024, 50 μM CK666, or 5 μM Latrunculin B for 5 minutes to allow for the drugs to inhibit. 100 uL of drugged 400 nM locking strand was then added for 10 minutes to allow for locking to take place and capture transient signal. Final images were taken in TRITC and RICM.

#### 3.12. Kinase Inhibitors

100 μL of CAR T cells were pretreated in incubator with either 1 μL untreated DMSO or 1 μL of a 100x drug stock to a final concentration of 50 nM PP2, 10 μM Zap180013, or 50 nM 17β-Hydroxywortmannin for 1 h. Cells were then allowed to spread on the tension probe surfaces for 10 minutes before adding 100 μL drugged 400 nM locking strand and allowing locking to occur for 10 min to capture signal accumulation. Final images were taken in TRITC and RICM.

#### 3.13. Dasatinib Treatment

All dasatinib-treated cells were pretreated in 0.25% DMSO or 1000 nM, 100 nM or 10 nM dasatinib drug in complete X-Vivo for 18-24 h.

For tension experiments, cells were spun down and resuspended in HBSS with the same concentration of dasatinib. Cells were then allowed to spread on the tension probe surfaces for 10 minutes before adding 100 μL drugged 400 nM locking strand and allowing locking to occur for 10 min to capture signal accumulation. Final images were taken in TRITC and RICM.

For cytotoxicity assays, cells were resuspended in fresh X-vivo. T cells and CAR T cells were co-cultured with 100,000 VPD-450 prestained CD19^+^ RS4;11s at 0.1:1, 0.2:1 and 0.3:1 effector to target ratios and dosed with DMSO, 1000 nM, 100 nM or 10 nM dasatinib drug in a U bottom 96 well plate for 24 hours at 37°C with 5% CO2.

The cells were stained with eFluor780 and APC Annexin V to determine live and dead cell populations. Samples were washed with 150 uL PBS, spun down at 320 × *g* for 5 min, then incubated with APC Annexin V and eFluor780 in Annexin V binding buffer for 30 minutes on ice. Cells that were VPD450^+^ eFluor780^−^ and APC Annexin V^−^ (SI Figure 5) were enumerated as the live cell percentage. At least 15,000 events were analyzed per condition in technical duplicates.

CAR-Specific Lysis was calculated by:

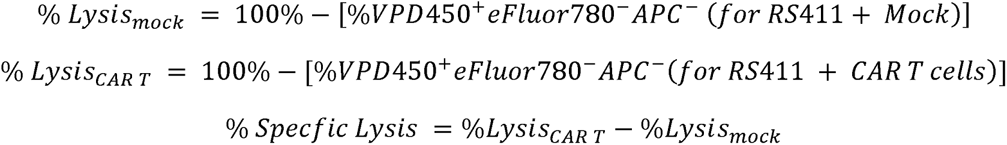

#### 3.14. Generating Alternative CAR Constructs

Additional CAR constructs were generated by altering the cDNA sequence from the original CD28-based CD19-CAR. In the CD19-41BB-CD3ζ CAR (41BB), the costimulatory domain was replaced using the intracellular domain from 4-1BB (CD137). The amino acid sequence for 4-1BB was obtained from UniProt and used to generate a codon optimized cDNA sequence which was then subcloned into the original vector. The CD19-CD28-CD3ζXXX CAR (ITAM K/O) was generated by substituting the two tyrosine residues within each CD3ζ ITAM with two phenylalanine residues, thus a total of six tyrosine residues were replaced for the three CD3ζ ITAM domains. The CD19 non-signaling CAR (nsCAR) was created by removing the CD28 co-stimulatory and intracellular CD3ζ domains from the original CAR. The CD28 transmembrane domain was retained to enable cell surface expression of the nsCAR. Finally, the CD33-CAR was generated by cloning a codon optimized HuM195 scFv cDNA sequence into the original second-generation CAR backbone. Cloned vectors were sent to Vectorbuilder for lentiviral vector production as described in section 3.1.

## Acknowledgments

This work was supported by the National Institutes of Health (NIH) grants through NIH NIAID R01 AI172452, NIGMS RM1 GM145394, and NIGMS R01 GM160947. We thank Brandon Fanelli for cloning plasmids, and Maia Vierengel and Brian Evavold for valuable discussions. Research reported in this publication was supported in part by the Pediatrics/Winship Flow Cytometry Core of Winship Cancer Institute of Emory University, Children’s Healthcare of Atlanta and NIH/NCI under award number P30CA138292. The content is solely the responsibility of the authors and does not necessarily represent the official views of the National Institutes of Health.

